# TAOK2 is an ER-localized Kinase that Catalyzes the Dynamic Tethering of ER to Microtubules

**DOI:** 10.1101/2021.04.22.440958

**Authors:** Kimya Nourbakhsh, Amy A. Ferreccio, Matthew J. Bernard, Smita Yadav

**Affiliations:** Department of Pharmacology, University of Washington, Seattle, WA 98195

## Abstract

The endoplasmic reticulum (ER) depends on extensive association with the microtubule cytoskeleton for its structure, function and mitotic inheritance. The identity of molecular tethers that mediate ER-microtubule coupling, and mechanisms through which dynamic tethering is regulated are poorly understood. Here, we identify, Thousand And One amino acid Kinase 2 (TAOK2) as a pleiotropic protein kinase that mediates tethering of ER to microtubules. We show that TAOK2 is a unique multipass membrane spanning serine/threonine kinase localized in distinct ER domains via four transmembrane and amphipathic helices. Using *in vitro* and cellular assays, we find that TAOK2 directly binds microtubules with high affinity. We define the minimal TAOK2 determinants that induce ER-microtubule tethering, and delineate the mechanism for its autoregulation. While ER membrane dynamics are increased in TAOK2 knockout cells, the movement of ER along growing microtubule plus-ends is disrupted. We show that ER-microtubule tethering is tightly regulated by catalytic activity of TAOK2 in both interphase and mitotic cells, perturbation of which leads to profound defects in ER morphology and cell division. Our study identifies TAOK2 as an ER-microtubule tether, and reveals a kinase-regulated mechanism for control of ER dynamics critical for cell growth and division.

## INTRODUCTION

The endoplasmic reticulum is an expansive and complex membranous cellular organelle. Composed of a continuous interconnected web of membrane sheets and tubules, the ER has distinct domains; namely the nuclear envelope, the rough ER sheets and smooth peripheral ER tubules (Voeltz et al., 2002). In addition, the ER makes specialized membrane contact sites with other organelles and the plasma membrane (Scorrano et al., 2019; Wu et al., 2018). Each of these domains are thought to be structurally discrete regions of the ER serving specialized physiological functions. This vast network of ER membranes relies on the microtubule cytoskeleton not only for structurally supporting its intricate shape and functional domains, but also for its motility and remodeling in response to stimuli. In animal cells, ER tubules align along microtubules (Terasaki et al., 1986). Disruption of microtubules by depolymerization agent nocodazole collapses the reticulated ER network into primarily ER sheets around the nucleus (Terasaki and Reese, 1994). In addition to its structural dependence, movement of ER membrane tubules occurs on microtubule tracks. Stabilization of microtubules by the drug taxol prevents new microtubule growth, and also inhibits ER tubule extension (Terasaki and Reese, 1994). Motor mediated ER ‘sliding’ movement occurs on stable acetylated microtubules (Friedman et al., 2010). Motor independent ER tubule extension along growing microtubule plus-ends is also dependent on microtubules, and is carried out by the ‘tip attachment complex’ composed of ER protein Stim1 and microtubule plus-end protein EB1 (Waterman-Storer and Salmon, 1998).

Additionally, membrane contact sites between ER and organelles appear to be supported by microtubules. For example, ER-mitochondria contact sites are preferentially aligned with acetylated microtubules (Friedman et al., 2010). Endosome maturation occurs at junctions where ER-endosome contact sites and microtubules converge (Wu and Voeltz, 2021). Store-operated calcium entry occurs through the Stim1-Orai channels at ER-plasma membrane contact sites (Grigoriev et al., 2008). Stim1 interaction with microtubules plays a facilitative role in organizing Stim1 for optimal Ca2+ sensing through its interaction with Orai channels (Smyth et al., 2007). In geometrically complex cells such as neurons, the ER is dependent on microtubules for its distribution throughout the fine processes such as dendrites, dendritic spines and axons. Presence of fine caliber ER tubules in axons is critical for neuronal polarity, and is dependent on ER membrane interactions with microtubules (Farías et al., 2019). On the other hand, unlike elongated ER in axons and dendrites, which is closely associated with microtubules, ER organization at dendritic branch points and dendritic spines is structurally complex and exhibits decreased microtubule association (Cui-Wang et al., 2012). Bidirectional regulation of ER-microtubule tethering, therefore not only influences ER morphology but can also dictate cell shape and function.

Identity of molecular tethers that mediate ER-microtubule coupling, and mechanisms through which tethering is physiologically regulated are not well understood. Studying ER-microtubule association and its feedback control is challenging, owing to the considerably expansive morphology of the ER, one that exhibits extreme overlap with the microtubule cytoskeleton. The nanoscale organization of sites at which ER is coupled with microtubules and its molecular composition is unclear. Constantly in flux, ER membranes undergo a range of dynamic and structural changes including membrane extension, retraction, three-way junction formation and tubule fusion (Pendin et al., 2011). Cell division brings about extensive ER remodeling, causing the ER membranes to coalesce around the spindle poles, but remain largely absent from the mitotic microtubule spindle at metaphase (Jongsma et al., 2015); (Smyth et al., 2015). Before insight into ER-microtubule dynamics and its physiological regulation can be gained, the mechanisms through which molecular tethers drive ER-microtubule association must be understood. Here, we investigate ER-microtubule association in interphase and mitotic human cells, and identify TAOK2 as an ER-localized multifunctional protein kinase that serves as a molecular tether linking ER to microtubules. An ER-microtubule tether should in principle be ER localized and be capable of direct and strong binding to microtubules. The tether would likely be under physiological regulation such that ER-microtubule binding can not only be ectopically induced, but also importantly turned off. Loss of molecular tether should disrupt ER association with microtubules. Our study demonstrates that the kinase TAOK2 meets all the above criteria of a *bona fide* ER-microtubule tethering molecule.

Thousand And One amino acid (TAO) kinases are ubiquitously expressed serine/threonine protein kinases belonging to the Ste20 kinase family (Chen et al., 1999; Manning et al., 2002). While there is only one *Tao* kinase encoding gene in invertebrates, three distinct *TAOK* genes are expressed in human (Chen et al., 2003; Manning et al., 2002). The encoded kinases TAOK1, TAOK2 and TAOK3 share a highly conserved N-terminal kinase domain, followed by distinct C-terminal domains (Chen et al., 2003; Manning et al., 2002). TAO kinases were originally identified as stress-sensitive kinases that activate the p38 kinase cascades through activation of MEK kinases (Chen et al., 1999). Among the TAO family of protein kinases, TAOK2 coordinates several aspects of neuronal development and function. TAOK2 is highly expressed during neuronal development and is important for basal dendrite formation as well as for axon elongation in cortical neurons (de Anda et al., 2012). TAOK2 is enriched in dendritic spines and is essential for their formation and stability (de Anda et al., 2012; Ultanir et al., 2014; Yadav et al., 2017). TAOK2 knockout mice exhibit cognitive and social-behavioral deficits, and show structural changes in brain size (Richter et al., 2018). Recently, mutations is TAOK2 have been associated with autism spectrum disorder, with several mutations present outside the kinase domain of the protein (Richter et al., 2018). Despite its clear relevance to human development and its disease association, little is known about the molecular and cellular functions of TAOK2 kinase.

In our study, we demonstrate that TAOK2 is a pleiotropic protein, with distinct catalytic kinase and ER-microtubule tethering functions, mediated through defined and dissociable domains. TAOK2 is enriched at junctions where the ER membrane makes contacts with the microtubule cytoskeleton. We show that TAOK2 is embedded in the ER membrane through transmembrane helices, and an amphipathic region that limits its localization to discrete subdomains of the ER. The cytoplasm facing ER-anchored C-terminal tail of TAOK2 directly binds microtubules with high specificity, and is essential for tethering of the ER membranes to microtubules. The ER and microtubule binding domains together, a remarkably small region of 100 residues, are sufficient for inducing ectopic tethering of ER to microtubules. Aberrant tethering by this minimal tether abolishes ER dynamics and mitotic ER remodeling. Importantly, we show that the tethering function of TAOK2 is negatively regulated by its kinase activity. During mitosis, we find that TAOK2 is highly activated, and inhibition of its catalytic function prevents ER disengagement from the mitotic spindle causing profound mitotic defects. This study identifies TAOK2 as an ER protein kinase and elucidates a hitherto unknown autoregulated mechanism for ER-microtubule tethering important for ER dynamics and mitotic segregation.

## RESULTS

### TAOK2 is a multi-transmembrane protein kinase that resides on the endoplasmic reticulum

To investigate the unique role that non-kinase domains of TAO family members might impart on their biological function, we performed bioinformatic analysis of secondary protein structure of TAO kinases. We found that TAOK2*α* (hereafter TAOK2) harbors unique hydrophobic regions in its C-terminal domain not present is its alternatively spliced isoform TAOK2*β*or paralogous genes TAOK1 and TAOK3. Sequence analysis of TAOK2 through the transmembrane helix prediction software TMHMM2.0 (Krogh et al., 2001) indicates that TAOK2 is a multipass membrane protein containing four transmembrane helices (Figures 1A and 1B). Additional predicted region of hydrophobicity following the fourth transmembrane domain was analyzed using AMPHIPASEEK (Combet et al., 2000) and HeliQuest (Gautier et al., 2008). This analysis revealed an amphipathic helical (AH) region with a sharply defined hydrophobic face (hydrophobicity <H>=0.75, hydrophobic moment <MM>=0.4) and a polar face rich in positively charged residues (net charge z=4) (Figure 1A inset).

**Figure 1.**
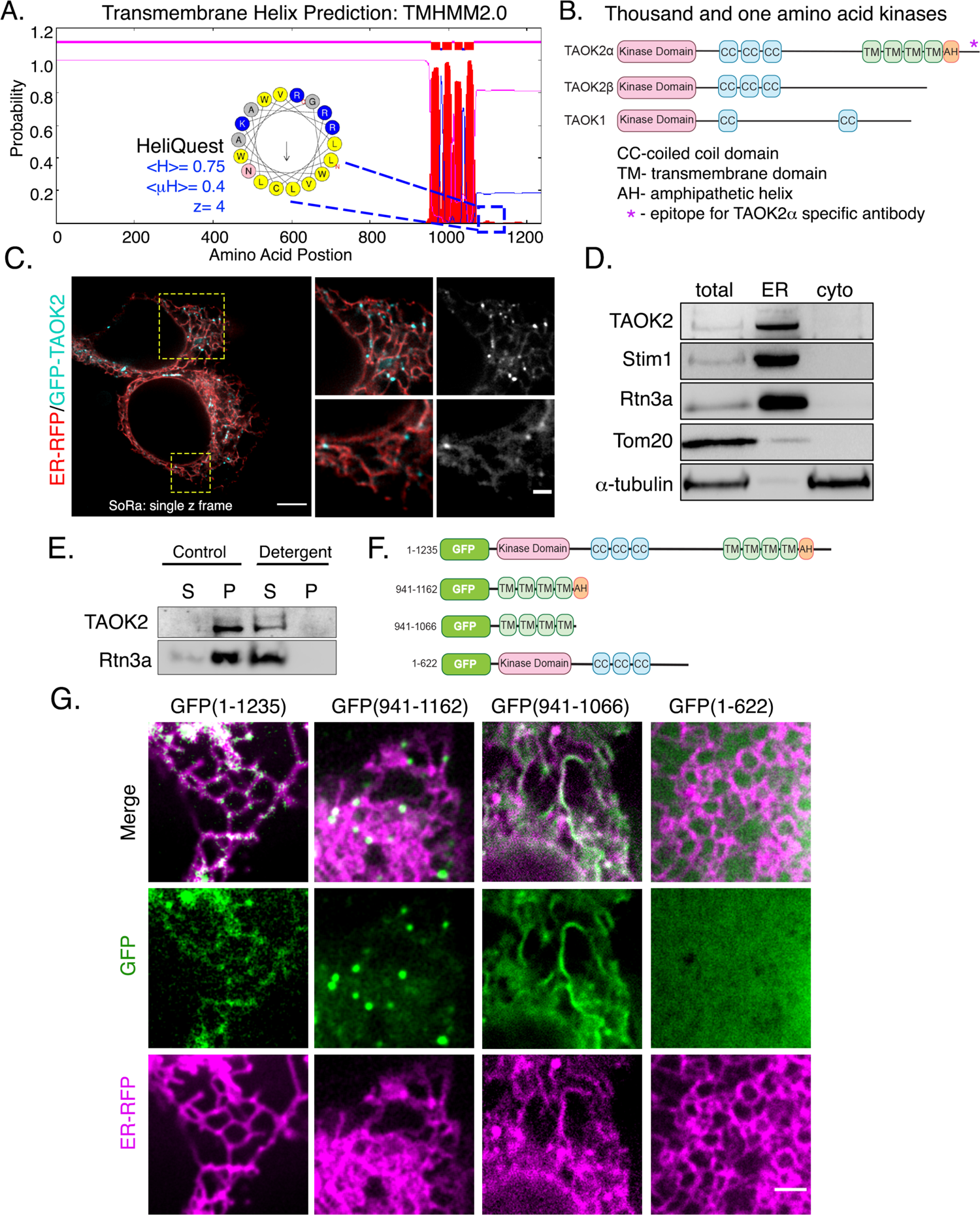
TAOK2 is an ER protein kinase that localizes to the ER membrane through four transmembrane domains and an amphipathic helix. (A) Transmembrane Hidden Markov Model (TMHMM v2.0) prediction plot shows the posterior probabilities (y axis) of inside/outside(magenta) / TM helix (red) along the length of TAOK2 sequence (x axis). Hydrophobic region following the 4^th^ predicted TMD was analyzed through AMPHIPASEEK which predicted residues 1146-1162 to be amphipathic. Charged (blue) and hydrophobic (yellow) residues are indicated in the inset. HeliQuest was used to calculate hydrophobicity <H>, hydrophobic moment <MH> and net charge z. (B) Schematic representation of TAOK2 isoforms α, β and TAOK1. The coiled coils (CC), four transmembrane domains (TM) and amphipathic helix (AH) predicted in TAOK2 are depicted. The unique C-terminal tail region of TAOK2 marked by the asterisk indicates the epitope used to generate the TAOK2α specific antibody. (C) Super resolution (SoRa: super resolution by optical reassignment) images of HEK293T cells expressing GFP-TAOK2 (cyan) and ER marker ER-mRFP (red), zoomed in sections are depicted in yellow boxes. Scale bar bottom left is 5mm, and bottom right is 0.5mm. (D) Western blot of cell homogenate fractionated into ER membrane and cytosol probed with antibodies against TAOK2, known ER membrane proteins Stim1 and Rtn3, mitochondrial protein Tom20 and tubulin. (E) Western blot of cell homogenate fractionated into membrane pellet (P) and cytosolic supernatant (S) components, in the absence (control) or presence of detergent, probed with antibodies against TAOK2 and ER protein Rtn3a. (F) Schematic shows the GFP-tagged TAOK2 deletion constructs used in Fig.1g. (G) Confocal images of HEK293T cells expressing distinct GFP-tagged TAOK2 deletion constructs (green) and ER-mRFP (magenta). Scale bar is 3mm.

To determine the cellular localization of the membrane spanning TAOK2 kinase, we generated a rabbit polyclonal antibody against the unique C-terminal tail (residues 1220-1235) of TAOK2*α*(Figure 1B). Antibody staining in HEK293T cells revealed that endogenous TAOK2 colocalized extensively with the ER protein calreticulin, as well as with the expressed ER marker EGFP-Sec22b (Figures S1A and S1B). We noted that TAOK2 exhibited a striking localization on subdomains of the ER membrane in distinct punctate pattern. Therefore, we generated a GFP-tagged TAOK2 construct and performed super resolution microscopy (super-resolution by optical reassignment using SoRa disk) to visualize the ER localization of TAOK2 at higher resolution (Figure 1C).

GFP-TAOK2 localized on the ER membrane and was present in discrete membrane subdomains (Supplementary Movie 1). To test biochemically whether TAOK2 is an ER membrane protein, we fractionated HEK293T cell homogenates into ER membrane fraction using differential centrifugation (Hoyer et al., 2018). TAOK2 was enriched in the ER membrane fraction along with other known ER membrane proteins such as Stim1 (Grigoriev et al., 2008) and Rtn3a (Hu et al., 2009). 97.6% of the total TAOK2 in the postnuclear homogenate was enriched in the ER fraction, as compared to 6.17% of tubulin (Figure 1D). On differential centrifugation, TAOK2 partitioned in the membrane fraction but not the cytoplasmic fraction, and treatment with detergent led to its release in the supernatant (Figure 1E). To identify the mechanism through which TAOK2 achieves its ER localization, we generated four GFP-tagged deletion constructs (Figure 1F).

Deletion construct lacking the transmembrane and amphipathic helices (1-622) was entirely cytosolic, while the four transmembrane helices alone (941-1066) were sufficient to target the kinase to the ER. A GFP-construct containing just the TAOK2 transmembrane domains localized uniformly throughout the ER, and was not restricted to ER subdomains. We found that the predicted amphipathic helical region together with the transmembrane domains (941-1162) is required for localization of TAOK2 to discrete ER subdomains, and captures the localization patters of the endogenous TAOK2 protein (Figure 1F). Therefore, using superresolution microcopy and biochemical assays, we show that TAOK2 is an ER resident protein kinase. Further, our data show that TAOK2 associates with the ER through its four transmembrane, while the amphipathic helices confer its localization to distinct punctate ER subdomains.

### TAOK2 associates directly with assembled microtubules via its conserved C-terminal tail

TAOK2 has previously been shown to colocalize with microtubules (Mitsopoulos et al., 2003), however mechanism underlying this observation is unknown. We found that overexpression of the C-terminal TAOK2 constructs led to bundling of microtubules, and these bundles were extensively decorated with GFP-TAOK2 constructs (Figure 2A and 2B). Expression of C-terminal GFP-tagged deletion constructs allowed us to map the microtubule-binding domain to 40 amino acids (1196-1235) in the extreme C-terminal tail of TAOK2 (Figure 2C). We next tested whether TAOK2 can bind microtubules directly using a biochemical binding assay with purified components. We bacterially purified GST tagged TAOK2 C-terminal tail protein (residues 1187-1235) (Figure S2A). An *in vitro* microtubule binding assay was used to test whether purified GST-TAOK2-C could bind microtubules polymerized from purified tubulin protein. GST-TAOK2-C pelleted specifically with polymerized MTs on centrifugation, while the control GST protein remained in the supernatant (Figure 2D and 2E), suggesting TAOK2 can directly associate with microtubules via its C-terminal tail. Further, the affinity of the TAOK2-microtubules interaction was assessed by determining the fraction of microtubule bound TAOK2-C at increasing concentration of tubulin. We found that the C terminal tail of TAOK2 associates with microtubules with a K_D_ of 0.67 ± 0.19μM (Figure 2F and 2G). These results show that TAOK2 can directly bind microtubules through its cytoplasm facing C-terminal tail.

**Figure 2.**
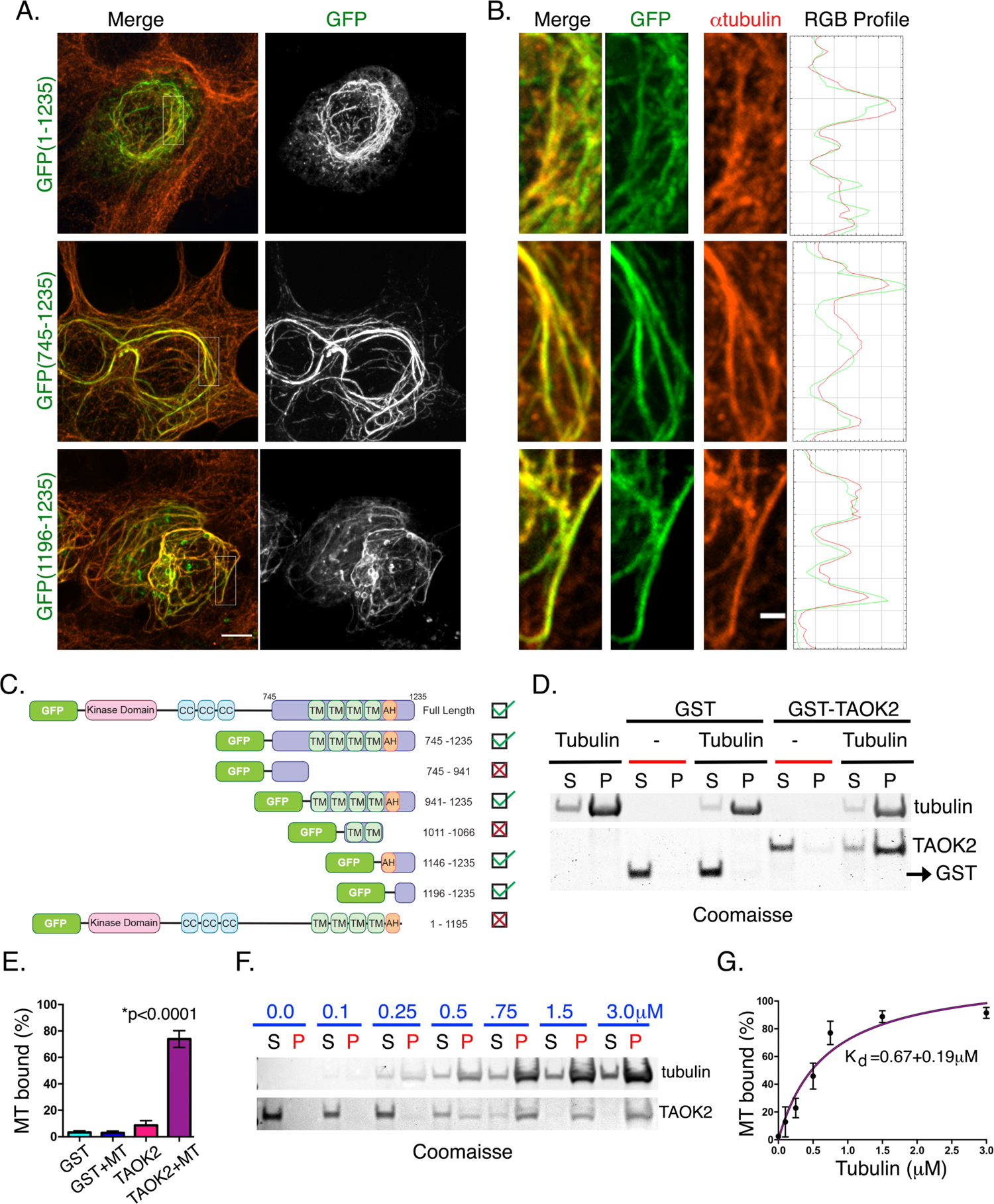
Direct binding of TAOK2 to microtubules through its C-terminal tail. (A) Confocal images of HEK293T cells expressing the indicated GFP-tagged TAOK2 construct immunostained for α-tubulin. Scale bar is 3mm. (B) Zoomed in images showing the overlap of GFP-tagged TAOK2 with microtubules immunostained for α-tubulin. Scale bar is 1mm. RGB profile of fluorescence intensity peaks of TAOK2 (green) and α tubulin (red). (C) Schematic representation of GFP-tagged deletion constructs used to map the microtubule binding domain. Coiled coil (CC), transmembrane (TMD); amphipathic helix (AH) domains are marked. Observed microtubule localization is indicated accordingly in the checkbox. (D) Coomassie stained SDS-page gel shows co-sedimentation of indicated proteins with polymerized tubulin. Microtubule binding assay was performed with 5μg of GST or GST-TAOK2-C (1187-1235) protein in the presence of taxol polymerized microtubules. Binding is assessed by the fraction of protein pelleted with microtubules (P) while unbound protein is in supernatant (S). (E) Percent protein bound to microtubules is plotted for each protein as indicated. Error bars indicate standard error of mean, n=3, p<0.0001, one-way ANOVA. (F) Coomassie stained SDS-page gel shows the amount of purified GST-TAOK2-(1187-1235) unbound (S) or co-sedimented with polymerized microtubules (P) with increasing concentrations of tubulin as indicated. (G) Strength of microtubule association was determined from dissociation constant K_D_ derived by Michaelis-Menten equation. Error bars indicate standard error of mean, n=3 replicates.

### ER membrane tethering to the microtubule cytoskeleton is mediated by TAOK2

Our finding that TAOK2 is an ER kinase with the ability to directly bind microtubules led us to hypothesize that this enzyme can act as a molecular tether linking the ER membrane to microtubules (Figure 3A). To evaluate this possibility, we performed live cell imaging with simultaneous visualization of the ER membrane and microtubule cytoskeleton. Cells transfected with full length GFP-TAOK2 and ER-mRFP were incubated with a cell permeable microtubule binding dye, and three color time-lapse confocal microscopy was performed (Supplementary Movie 2). Indeed, 87% of GFP-TAOK2 punctae localized at the points of contact between the ER and the microtubule cytoskeleton (Figure 3A-3C). These punctae tracked with the movement of the ER membrane on microtubules (Supplementary Movie 2). In order to ascertain the domains within TAOK2 that conferred the tethering function of TAOK2, we generated several constructs that retained the ER localization and microtubule binding elements. The first included the transmembrane domain (TMD), the amphipathic helix (AH) and microtubule binding domain, but lacked the N-terminal kinase and coiled coil domains (Figure 3D, top row). We found that expression of this construct led to aberrant over-tethering perturbing ER morphology, such that the ER appeared bundled alongside perinuclear microtubule cables. Next, we expressed a minimal-tether construct containing just the AH linked to the microtubule binding domain (1146-1235). Expression of the ‘mini-tether’ led to a complete collapse of the ER on to the microtubule cytoskeleton, where the ER membrane was entirely conformed to the shape of the microtubule cytoskeleton (Figure 3D, center row). Live imaging revealed that aberrant tethering induced by the minimal-tether abolishes ER membrane dynamics due to its collapse on microtubules (Supplementary Movie 3). As anticipated, the TMD and the AH domain without the microtubule binding domain failed to induce tethering, and the ER retained its lace-like morphology occupying the entire cell area (Figure 3D, bottom row). These data together define the mechanisms through which TAOK2 mediates its tethering function, and delineate the minimal elements comprising the ER-microtubule tether. Importantly, these findings also imply that additional domains of TAOK2 outside the tethering elements regulate the coupling dynamics of the ER to the microtubule cytoskeleton.

**Figure 3.**
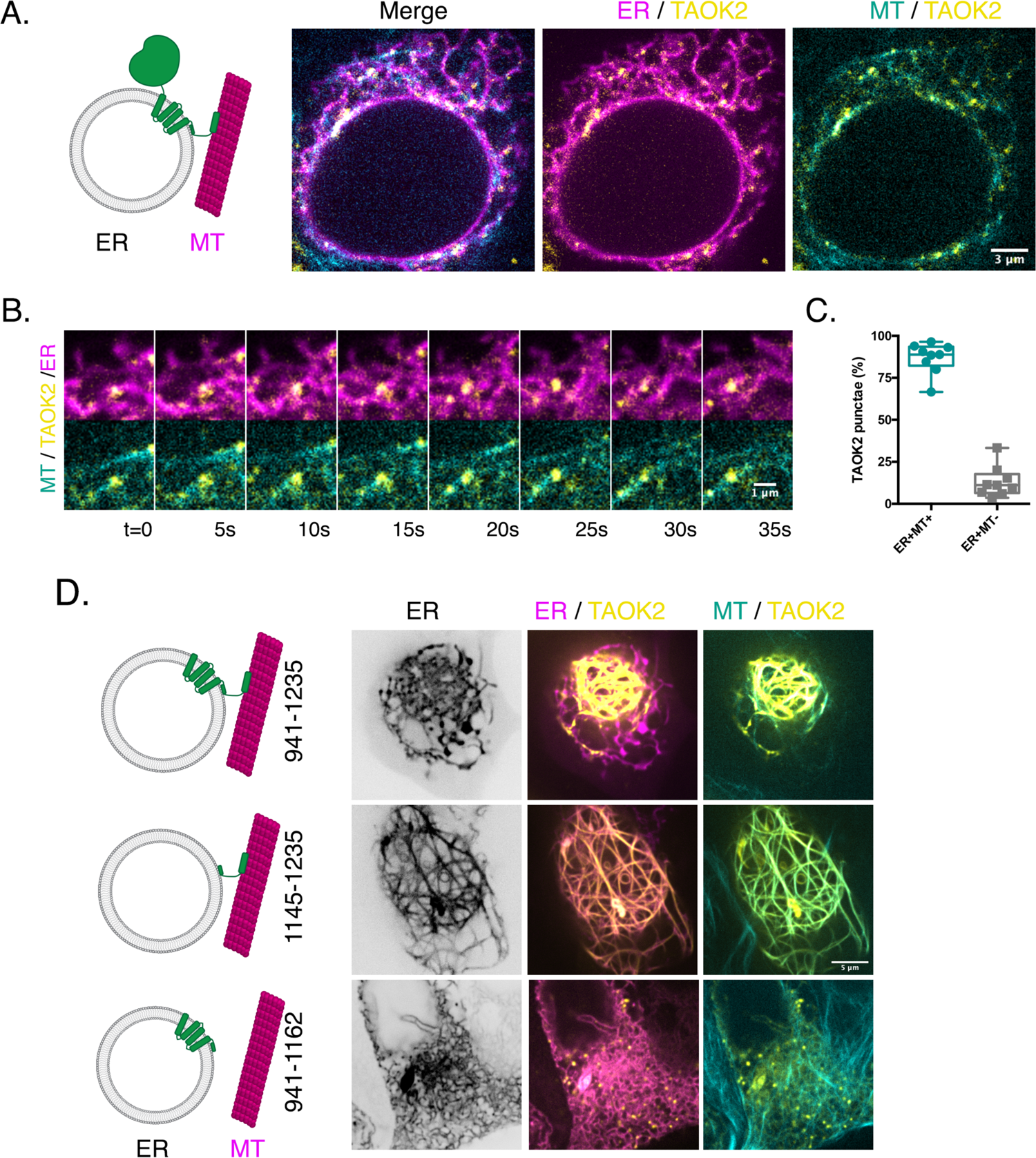
TAOK2 is an ER-microtubule tethering protein kinase. (A) TAOK2 (green) schematically depicted as a multipass transmembrane protein on the ER (grey) has an N-terminal cytoplasm facing kinase domain and a C-terminal tail that directly binds microtubules (magenta). Confocal images of HEK293T cells expressing GFP-TAOK2 (yellow), ER-mRFP (magenta) and live-stained with microtubule dye (cyan). Scale bar is 3μm. (B) Montage of confocal time lapse images shows TAOK2 (yellow) punctae colocalized with both ER membrane (magenta) and microtubules (cyan). Scale bar is 1mm. (C) Percent TAOK2 puncta colocalized with ER and microtubules is plotted. Values indicate mean, n=10 cells and error bars indicate S.E.M. (D) Schematic on the left depicts topology of TAOK2 deletion constructs corresponding to representative confocal images of HEK293T cells expressing the indicated TAOK2 construct (yellow), ER-mRFP (magenta) and microtubules (cyan). Scale bar is 5mm.

### TAOK2 is required for ER microtubule plus end motility but not motor mediated movement

The ER membranes utilize the microtubule cytoskeleton as tracks for movement^13-15^. Distinct mechanisms of ER motility have been defined; ‘sliding’ movements are kinesin-based rapid movement, and ‘tip-attachment complex’ movements occur when the ER attaches to the MT plus ends and tracks along the growing MT (Friedman et al., 2010; Waterman-Storer and Salmon, 1998). To determine the functional consequences of TAOK2 depletion on ER motility and ER-MT movement, we generated two independent TAOK2 knockout (KO) HEK293T cell lines using CRISPR/Cas9 mediated gene editing. Genetic knockout and loss of protein was validated using genome sequencing and western blot respectively (Figure 4A, Figure S2B and S2C). Wildtype and TAOK2 KO cells were transfected with the ER marker EGFP-Sec22b and time-lapse confocal microscopy allowed us to visualize ER dynamics. To test whether loss of TAOK2 mediated tethering would impact overall motility of the ER membranes, we measured ER membrane movement over time. ER motility in TAOK2 KO cells was significantly increased compared to WT cells (Figure 4B). Average normalized pixel differences over time allowed us to calculate the ER motility index which increased from 0.287±0.012 in WT cells to 0.351±0.017 (n=9, p=0.0098) in TAOK2 KO cells (Figure 4C). To further investigate how absence of TAOK2 mediated tethering might increase ER motility, we performed simultaneous confocal imaging of ER (EGFP-Sec22b) and microtubule end binding protein mCherry-EB3. We found that in WT cells both TAC movements on microtubule plus end, as well as sliding ER movements were observed, however, in TAOK2 KO cells almost all observed movements were mediated by sliding movements. Thus, loss of TAOK2 disrupts TAC movements (Figure 4D), while motor mediated ER sliding movements are maintained (Figure 4E). Next, we determined whether absence of TAOK2 would impact microtubule dynamics. We found that microtubule growth assessed by measuring EB3 velocity was slightly increased in TAOK2 KO cells (Figure 4F). Further, in TAOK2 knockout cells, EB3 comet tracks were less directed exhibiting increased curvature and paused more frequently compared to control WT cells (Figure 4F-4G, Supplementary Movie 4). Thus, our analyses of the TAOK2 knockout reveal an important role of TAOK2 in regulating the dynamics of ER-microtubule based movement, and suggest that TAOK2 mediated ER-MT tethering is essential for the structure and dynamics of ER membranes.

**Figure 4.**
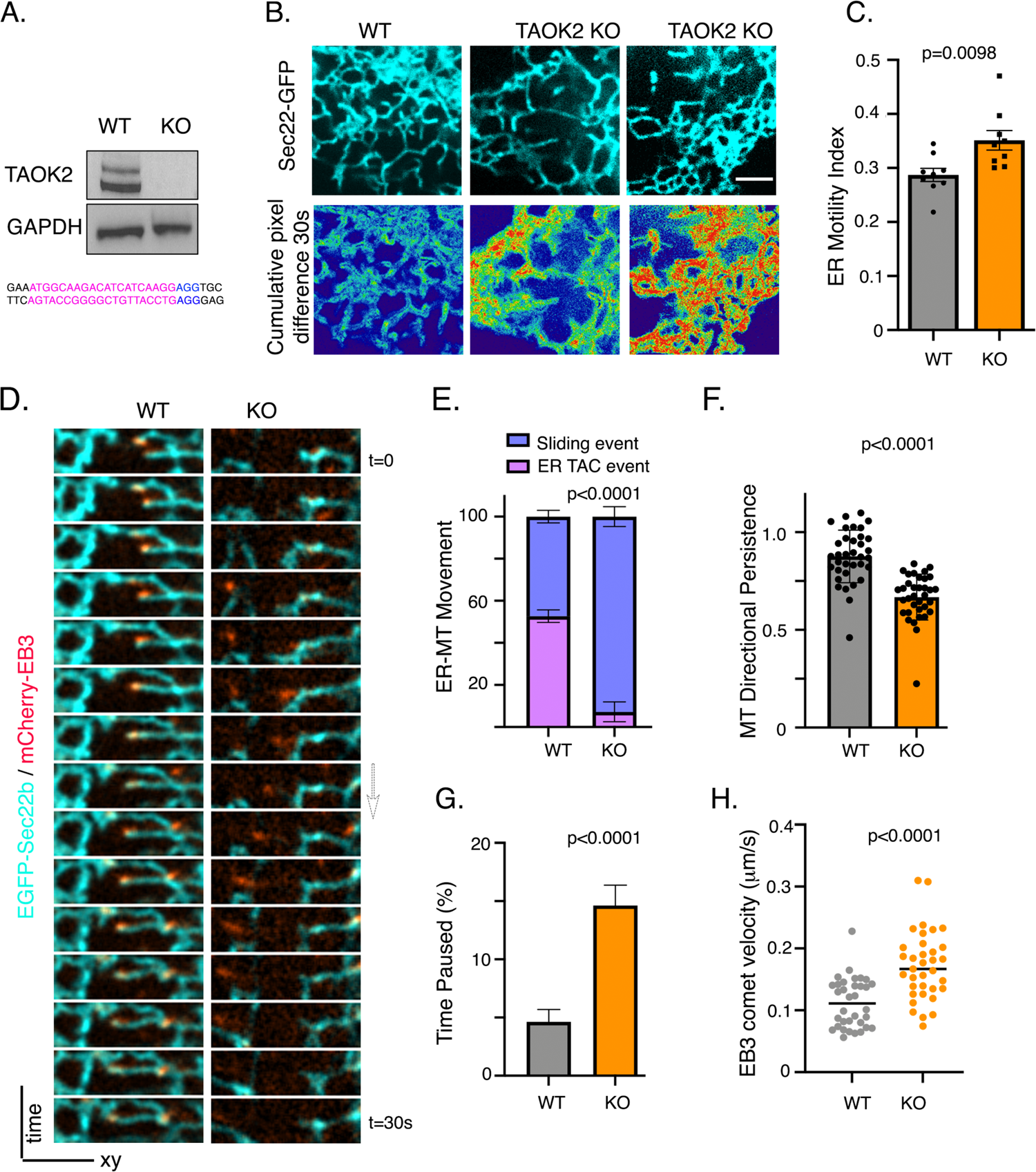
TAOK2 knockout disrupts ER-microtubule dynamics. (A) Western blot of lysate from wildtype and TAOK2 (knock out) KO cells generated using CRISPR/Cas9 mediated gene editing. Guide RNA sequences used for the knockout are shown. (B) Confocal images of peripheral ER in wildtype and TAOK2 KO cells expressing ER marker EGFP-Sec22b at a single time point (left column). Sum slice projection of cumulative pixel difference in successive frames over 30s time period (right), where images were acquired every 3sec. Fluorescence intensity is pseudocolor coded, red regions representing increased ER motility. Scale bar is 3mm. (C) ER motility index calculated by averaging the normalized pixel difference between successive time frame over 30s is plotted for wildtype and TAOK2 KO cells. Error bars indicate S.E.M, n=9 cells, t-test. (D) Montage of images acquired every 2s, of wildtype and TAOK2 KO cells expressing EGFP-Sec22b and mCherry-EB3 shows movement of ER membranes (cyan) associated with microtubule plus tips (red) in wildtype but not KO cells. (E) Time lapse confocal images of cells expressing EGFP-Sec22b and mCherry-EB3 were analyzed and ER membrane movements classified into slow MT plus tip associated-TAC (colocalized with EB3 comet) and fast motor driven sliding movements (lacking association with EB3 comets). Percent MT plus tip associated (TAC) events and sliding events is shown for WT and TAOK2 KO cells. Mean values are plotted, n=5 cells for each condition. (F) Microtubule directional persistence, calculated as a fraction of perpendicular distance between start/end points and the length of the actual path taken is plotted for wildtype and TAOK2 knockout cells. Values indicate mean ± S.E.M., n=9 cells with at least 5 comet paths measured per cell, two tailed t-test. (G) Percent total time spent by EB3 comet pausing (no growth) is plotted for wildtype and TAOK2 KO cells. Values indicate mean ± S.E.M., n=9 cells with at least 5 comet paths measured per cell, two tailed t-test. (H) EB3 comet tracks generated using the Manual tracking function in Fiji were used to measure EB3 comet velocity by dividing the total distance traveled over time. Mean values for wildtype and TAOK2 KO cells are plotted, error bars indicate S.E.M, n=9 cells with 5 comets per cell, two tailed t-test.

### Aberrant TAOK2 tethering disrupts ER restructuring during mitosis

The endoplasmic reticulum undergoes dynamic restructuring during cell division (Carlton et al., 2020). During metaphase, as the mitotic spindle aligns the chromosomes at the metaphase plate, the ER is anchored at each end to the spindle poles but largely absent from chromosomes and the area between the spindle poles. Defects in ER clearance from the chromosomes and spindle elicit mitotic defects (Schlaitz et al., 2013). Given the extensive association of ER with MT during cell division, we investigated the role of TAOK2 in ER restructuring during mitosis. Immunofluorescence using the C-term TAOK2 antibody revealed that TAOK2 was present as discrete puncta throughout the ER, the spindle pole, and on the spindle MT during mitosis (Figure 5A). We performed superresolution confocal microscopy to visualize GFP-TAOK2 localization in mitotic cells. We found that GFP-TAOK2 was present in close apposition to the spindle poles at sites where the ER membranes converged. TAOK2 localized at discrete punctate sites throughout the curvilinear peripheral ER surrounding the mitotic cell, as well as at the points of contact between ER membrane and the mitotic spindle (Figure 5B, bottom row). To test, whether TAOK2 was important for ER structural remodeling during cell division, we imaged WT and TAOK2-KO HEK293T mitotic cells expressing EGFP-Sec22b and mCherry-EB3 to visualize ER membranes and mitotic spindle. WT cells exhibited the characteristic ER morphology showing accumulation at spindle poles and fenestrated curvilinear peripheral ER. In contrast, TAOK2-KO cells had an abnormal morphology with increased peripheral curvilinear ER membranes, and concomitant decrease in association with spindle poles (Figure 5C and 5D). To determine if this aberrant ER morphology affected cell division, we immunostained WT and TAOK2 KO cells with tubulin and DAPI. While 95.6% WT cells showed normal bipolar spindle, only 44.8% KO cells had a normal bipolar spindle. KO mitotic cells displayed a chromosomal misalignment defect, where 39.3% KO cells had a bipolar spindle with misaligned chromosomes, and 15.8% KO cells had multipolar spindle (Figure 5E). While loss of TAOK2 induced defects in ER association with mitotic spindle, we next queried what would be the consequence of unregulated over-tethering by TAOK2. Expression of the short TAOK2 lacking the N-terminal kinase and coiled coil domains, induced a dramatic collapse of the ER membranes on the mitotic spindle (Figure 5F). TAOK2 binding to microtubules created an extremely short and stable MT-ER bridge between the spindle poles, which we refer to as pseudomonopolar (90.5% mitotic cells). The chromosomes were displaced from the metaphase plate and instead formed a rosette around the ER-MT spindle (Figure 5F, top row). These cells failed to divide. Expression of the ‘mini-tether’ construct containing the AH and the microtubule binding domain, generated a similar phenotype of pseudomonopolar spindle (30.7% mitotic cells) where the ER remained attached to the microtubules. Additionally, cells expressing the minimal tether displayed aberrant spindles which did not appear to be focused on the spindle poles (61.5% mitotic cells). In all the cases, the chromosomes were displaced from the spindle and ER was tightly associated with the spindle microtubules. The number of normal bipolar cells decreased from 96.7% in cells expressing WT-TAOK2 to 1.96% and 3.93% in mitotic cells expressing the tether constructs TAOK2 (941-1235) and TAOK2 (1146-1235), respectively. However, expression of a TAOK2 construct lacking the microtubule binding domain TAOK2 (941-1162), had normal ER morphology during mitosis, and 92.2% mitotic cells had a bipolar normal spindle (Figure 5F and 5G). These data collectively show that downregulation of TAOK2 mediated ER-MT tethering during cell division is required for the disengagement of ER from the mitotic spindle and its segregation into daughter cells.

**Figure 5.**
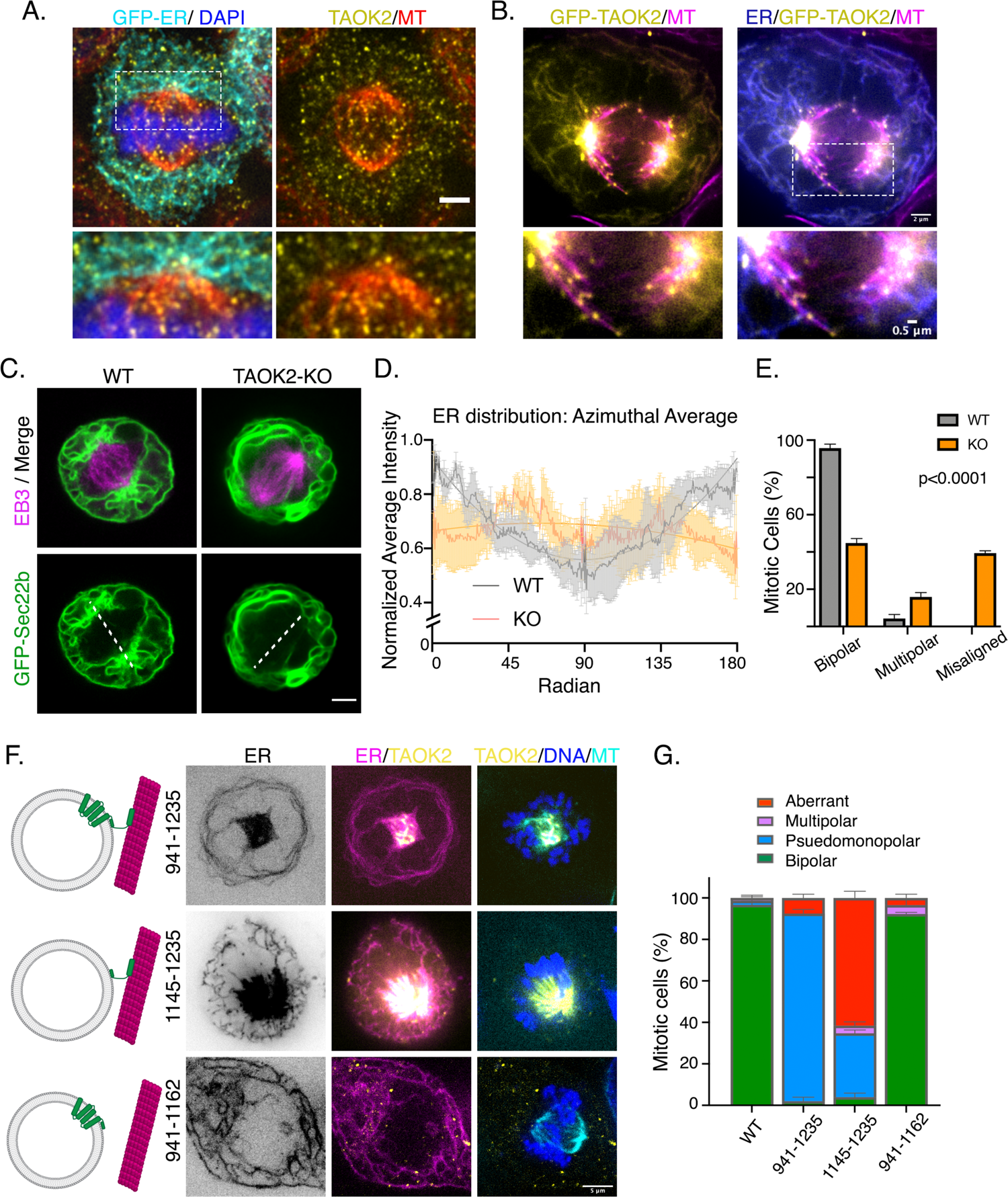
TAOK2 localizes to the mitotic spindle and regulates mitotic ER remodeling. (A) TAOK2 antibody staining shows localization of endogenous TAOK2 (yellow) on the ER (cyan) and mitotic spindle microtubules (red) in mitotic cells. (B) Superresolution confocal image of mitotic cell expressing GFP-TAOK2 (yellow) and ER-mRFP (blue) and stained with microtubule dye (magenta), scale bar is 2mm (top right). Magnified view of the ER-MT tethering sites on the spindle is shown in the bottom row, scale bar is 0.5mm. (C) Confocal live cell images of WT and TAOK2 KO mitotic cells expressing EGFP-Sec22b (green) and mCherry-EB3 (magenta). Dashed white line drawn across the spindle poles highlights the difference in spindle pole associated ER fraction in WT and TAOK2 KO cells. Scale bar is 5mm. (D) ER distribution in mitotic cells is measured using Azimuthal Average of normalized fluorescence intensity in each hemisphere demarcated by the mitotic spindle (where 0 and 180 correspond to the spindle poles) is plotted for WT (gray) and TAOK2 KO (orange) cells. Average means are plotted, error bars are S.E.M., and n=3 cells. (E) Percent mitotic WT (grey) and TAOK2 KO (orange) cells that exhibit bipolar, multipolar or misaligned mitotic spindles are plotted. Values indicate mean ± S.E.M., n>45 mitotic cells from three different experiments, p<0.0001 one-way ANOVA. (F) Schematic (left) depicts topology of GFP-TAOK2 deletion constructs. Corresponding confocal images of mitotic HEK293T cells expressing the indicated TAOK2 construct (yellow), ER-mRFP (magenta), DNA (blue) and microtubules (cyan) are shown. Scale bar is 5mm. (G) Mitotic defects in cells expressing the indicated TAOK2 constructs is depicted as the percent of mitotic cells exhibiting normal bipolar or aberrant spindles. Values indicate mean ± S.E.M., n=52 cells from 3 different experiments.

### Catalytic autoregulation of TAOK2 mediated ER-MT coupling

What might be the mechanism through which TAOK2 regulates its function as an ER-MT tether? We tested whether the kinase activity of TAOK2 regulates its microtubule association. First, we introduced a kinase-dead mutation within the catalytic domain of TAOK2 at residue K57A, which disrupts the autophosphorylation at the critical residue S181 in the activation loop (Moore et al., 2000), and hence the catalytic activity of TAOK2 (Figure 6A). Comparative analysis of cells expressing ER-mRFP along with either GFP tagged-TAOK2 WT or TAOK2-K57A revealed that loss of kinase activity increased the association of TAOK2 with microtubules (Figure 6B). Colocalization analysis of GFP positive TAOK2 puncta with tubulin showed that association with microtubules increased from 87.86±2.17% for TAOK2-WT to 96.46±1.28% for TAOK2-K57A (Figure 6C). To test if increased association with microtubules resulted in stronger ER-MT tethering, we measured ER motility in cells expressing kinase dead TAOK2-K57A and ER marker EGFP-Sec22b. A substantial decrease in ER membrane motility was found in cells expressing TAOK2-K57A as opposed to TAOK2-WT (Figure 6D and Supplementary Movies 5-6). The ER motility index calculated from mean normalized pixel differences over time, was found to decrease from 0.28±0.006 in TAOK2-WT to 0.23±0.01 (n=10, p=0.0029) in K57-TAOK2 expressing cells (Figure 6E). Further, we found that TAOK2 kinase activity was important for microtubule growth rates determined by measuring EB3 comet velocity. EB3 velocity averaged at 0.35μm/s in TAOK2-WT expressing cells and was decreased significantly to 0.17μm/s in cells expressing kinase dead TAOK2 (Figure 6F and Supplementary movies 5-6). While directional persistence of MT growth was reduced, the frequency of MT comet pauses increased from 8.5% in control to 19.7% in K57A-TAOK2 expressing cells (Figure S3A and S3B). These data indicate that catalytic activity of TAOK2 regulates its ER-MT tethering function. While TAOK2 kinase activity negatively regulates its association with microtubules, it positively regulates microtubule growth and dynamics.

**Figure 6.**
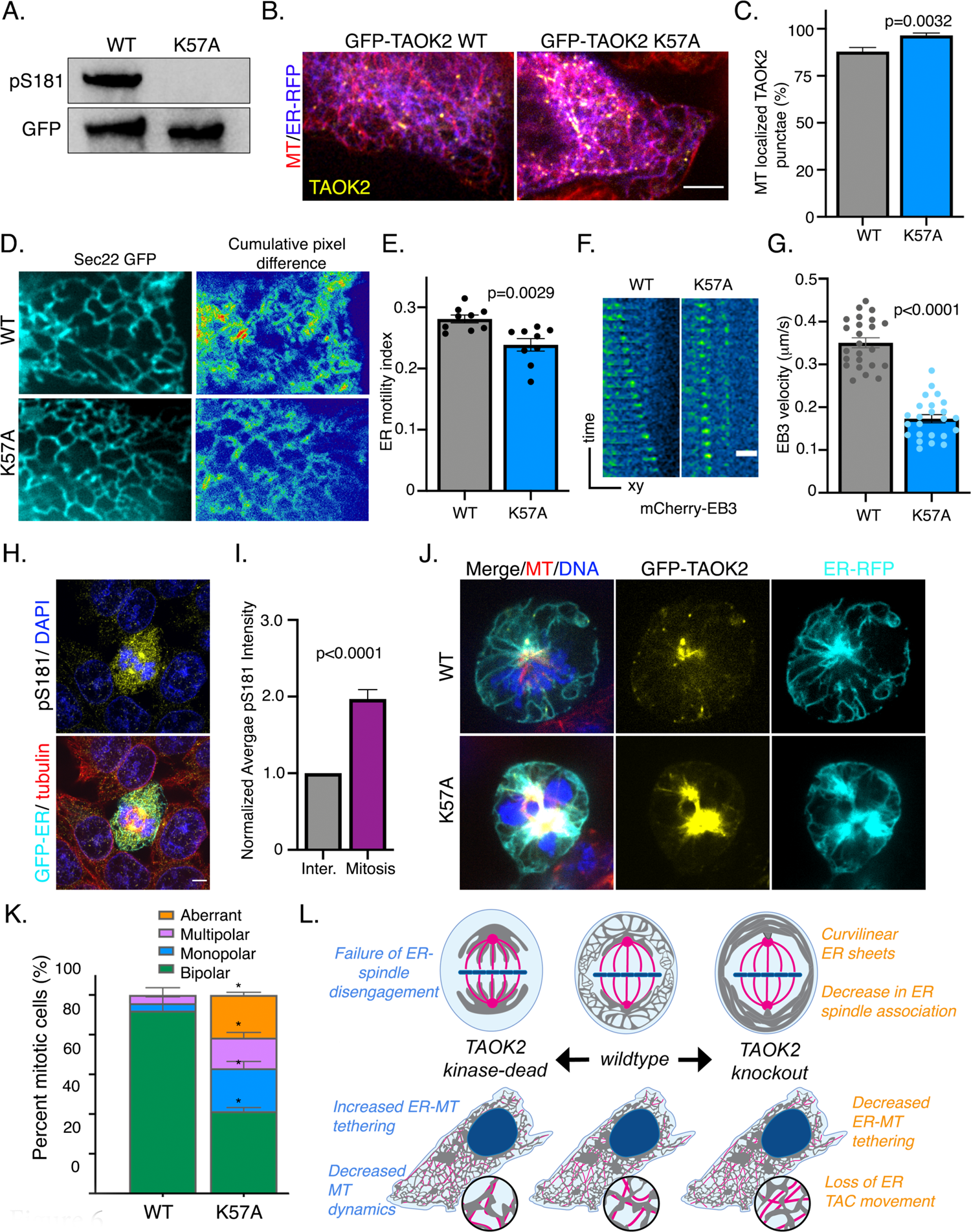
ER-microtubule tethering is regulated by catalytic activity of TAOK2. (A) Catalytic activity of GFP-TAOK2 WT and GFP-TAOK2 K57A was measured in an *in vitro* kinase reaction using autophosphorylation at S181 as the readout. Western blot probed with phospho-S181 antibody to measure kinase activity and anti-GFP represent the total amount of TAOK2. (B) Confocal images of cells expressing GFP-TAOK2 WT and GFP-TAOK2 K57A (yellow) along with ER-mRFP (blue) and live-stained with microtubule dye (red). Scale bar is 5mm. (C) Percent of GFP-TAOK2 WT and GFP-TAOK2 K57A puncta colocalized with microtubules is plotted. Values indicate mean, n=10 cells, error bars indicate S.E.M., p<0.005, two tailed t-test. (D) Confocal images of peripheral ER in cells expressing TAOK2 WT or TAOK2 K57A along with ER marker EGFP-Sec22b at a single time point (left column). Sum slice projection of cumulative pixel difference in successive frames over 30s, where images were acquired every 3s are shown. Fluorescence intensity is pseudocolor coded, red regions representing increased ER motility. Scale bar is 3mm. (E) ER motility index is calculated by averaging the normalized pixel difference between successive time frame over 30s. Mean ER motility for TAOK2 WT and TAOK2 K57A expressing cells is plotted. Error bars indicate S.E.M, n=10 cells, two tailed t-test. (F) Montage of images acquired every 2s of cells expressing TAOK2-WT or TAOK2 K57A along with mCherry-EB3 shows movement of EB3 comets (cyan) over time on y axis. Scale bar is 1mm. (G) EB3 comet tracks generated using the Manual Tracking function in Fiji were used to measure EB3 comet velocity by dividing the total distance travelled over time. Mean velocity values for cells expressing mCherry-EB3 along with either TAOK2 WT or TAOK2 K57A are plotted. Error bars indicate S.E.M, n=10 cells, two tailed t-test. (H) Confocal images of mitotic and interphase HEK293T cells stained with phospho-S181 and tubulin antibodies to label active TAOK2 and microtubules respectively. DNA stained with DAPI. (I) Average intensity of phospho-S181 staining in mitotic cells normalized by average interphase intensity is plotted. Error bars indicate S.E.M, n=10 cells, t-test with Welch correction. (J) Confocal images of mitotic cells expressing GFP-TAOK2 WT or GFP-TAOK2 K57A (yellow) along with ER-mRFP (cyan) was stained with DNA dye (blue) and microtubules dye (red). Scale bar is 5mm. (K) Mitotic defects in cells expressing GFP-TAOK2 WT or GFP-TAOK2 K57A are plotted as the percent of mitotic cells exhibiting normal bipolar or aberrant spindles. Values indicate mean ± S.E.M., n=50 cells from 3 different experiments. (L) Schematic representation of functional roles of TAOK2 in maintaining ER structure and its remodeling during mitosis. Distinct effects of TAOK2 depletion (TAOK2 knockout) compared to those due to kinase dysfunction (TAOK2 kinase dead) in interphase and mitotic cells are shown. The divergent effects of TAOK2 kinase dead mutation and TAOK2 knockout suggest pleiotropic roles of TAOK2 mediated by functional domains in addition to its kinase domain.

Next, we assessed whether TAOK2 catalytic activity was regulated during mitosis, and found that TAOK2 was highly activated during mitosis. Immunostaining interphase and mitotic cells with phospho-S181 antibody revealed a 2-fold increase in mitotic cells compared to interphase (Figure 6G). Western blot analysis showed high ratio of pS181-TAOK2/TAOK2 in lysates from synchronized mitotic cells compared to asynchronous cell lysate (Figure 6H). Therefore, we tested if perturbation of TAOK2 kinase activity would impact mitosis or ER segregation during cell cycle. We expressed either GFP tagged TAOK2-WT or TAOK2-K57A along with ER-mRFP, and then used a 405nm-DNA dye and 673nm-MT dye to enable four-color imaging of TAOK2, ER membrane, chromosomes and mitotic MT spindle. We found that while in TAOK2-WT expressing cells, both TAOK2 and the ER membranes were enriched at the spindle poles, and a majority of the ER membrane was dissociated from the spindle microtubules. However, in cells expressing TAOK2-K57A kinase dead mutant, TAOK2 association with the spindle MT was markedly increased and ER membranes were extensively associated with the mitotic spindle MT (Figure 6I). A failure of ER membranes to disengage from the spindle MT led to severe mitotic defects including monopolar (22%), multipolar (15%) and aberrant spindles (21%). Accordingly, the number of mitotic cells exhibiting a normal bipolar spindle deceased from 92% in cells expressing TAOK2-WT to about 41% in TAOK2-K57A expressing cells (Figure 6J).

These data demonstrate that the kinase activity of TAOK2 is important for dynamic regulation of ER-MT tethering during cell division, and perturbation of its catalytic activity leads to failure of ER segregation into daughter cells, ultimately inducing mitotic defects.

## DISCUSSION

Our study identifies TAOK2 as an ER resident kinase that functions as a molecular tether linking the ER membranes to microtubule cytoskeleton. TAOK2 is a large multifunctional protein of 1235 amino acids (Chen et al., 1999). At the molecular level, we show that distinct structural domains within TAOK2 confer its catalytic activity, ER localization and microtubule association. Importantly, we demonstrate that, while the ER-microtubule tethering function of TAOK2 is structurally dissociated from its kinase domain, the catalytic activity of TAOK2 negatively regulates ER-microtubule tethering.

Our results show that knockout of TAOK2 and kinase-dead TAOK2 expression have opposing effects on ER-microtubule tethering and dynamics in both interphase and mitotic cells. Thus, TAOK2 possesses the unique capacity to autoregulate ER-microtubule tethering through its kinase activity. We provide two key pieces of evidence in support of bidirectional autoregulation of tethering by TAOK2. First, we show that the kinase dead TAOK2 is a stronger ER-microtubule tether. Second, TAOK2 kinase activity is increased in mitosis, which correlates with a dramatic decrease in the tethering of ER membranes with the mitotic spindle microtubules. Perturbation of TAOK2 kinase activity in mitotic cells leads to failure in disengagement of ER from the spindle causing mitotic defects.

Little is known about upstream mechanisms that mediate TAOK2 activation. Early studies on the family of TAO kinases suggest that they might function as intermediate signaling kinases that link certain heterotrimeric G protein-coupled receptors to the p38 MAPK pathway. Among ligands that induce TAOK2 activation, nocodazole, sorbitol and the muscarinic agonist carbachol were found to increase TAOK2 activity from 1.5-3 fold on stimulation (Chen et al., 2003). Members of the TAO family can be activated by ATM kinase is response to genotoxic stress (Raman et al., 2007), however, whether TAOK2 is phosphorylated is unknown. TAOK2 is phosphorylated by the Hippo kinase mammalian homolog MST3 at residue T468 proximal to the kinase domain (Ultanir et al., 2014). Whether phosphorylation by MST3 increases TAOK2 catalytic activity has not been demonstrated. Further, in neurons, the secreted semaphorin molecule Sema3a has been shown to increase TAOK2 kinase activity in neurons through interaction with its receptor Neuropilin 1 (de Anda et al., 2012). Therefore, several independent signaling pathways might impinge on TAOK2 kinase to achieve distinct context-dependent cellular outcomes.

Our findings show that TAOK2 is a unique serine/threonine kinase, as no other multipass membrane serine/threonine kinase has been reported to date. Two other kinases with a single transmembrane domain reside in the ER membrane, Ire1(Cox et al., 1993; Mori et al., 1993) and PERK (Harding et al., 1999). Both Ire1 and PERK kinases have a cytosol facing kinase domain similar to TAOK2. The C-terminal tails of Ire1 and PERK project into the luminal domain where they have important functions associated with sensing ER stress (Ron and Walter, 2007). In contrast, the C-terminal tail of TAOK2 faces the cytoplasm and directly binds microtubules. The intrahelical loops between with the transmembrane helices are predicted to be extremely short (4-6 amino acids), and it is unlikely that TAOK2 serves a role as an ER stress sensor within the ER lumen. However, it is conceivable that TAOK2 is activated during ER stress indirectly through other ER stress sensors. Our data suggests that the punctate localization of TAOK2 within distinct ER subdomains is conferred by an amphipathic helical domain following the fourth transmembrane helix. Investigating the molecular identity of these ER subdomains in terms of its lipid and protein constituents would likely reveal important insight into TAOK2 regulation. Based on the findings of this study, we predict that increased activation in conditions of cellular stress might negatively regulate its function as an ER-MT tether, and will be an important area of future inquiry.

Identification of TAOK2 as a unique kinase that also functions as an ER-MT tether, adds to the short list of two other multifunctional ER enzymes, Spastin and Atlastin that bind microtubules. Hereditary spastic paraplegia protein, Spastin, is a microtubule severing AAA ATPase enzyme that localizes to the ER and remodels microtubules via its C terminus domain (Hazan et al., 1999; Roll-Mecak and Vale, 2008). The other is Atlastin, an ER localized GTPase required for ER tubule fusion (Orso et al., 2009). In addition to these enzymes, several other ER proteins associate with the microtubule cytoskeleton directly or indirectly, and are each likely to serve a particular physiological function through tethering (Wang et al., 2016). Perhaps the most well studied, Stim1, is an ER protein that interacts with microtubule growing tips indirectly by binding EB1 proteins (Grigoriev et al., 2008; Pavez et al., 2019). Climp63 is an ER protein thought to instruct the luminal width of ER sheets (Nikonov et al., 2007; Vedrenne et al., 2005) and can directly associate with microtubules (Klopfenstein et al., 1998). It is important to note, that unlike the abovementioned microtubule binding ER proteins, TAOK2 has the unique ability to bidirectionally autoregulate its microtubule association.

As a kinase highly expressed in neurons, and one critical for neurodevelopment, the role of TAOK2 as an autoregulated ER-MT tether is likely to serve specific physiological functions in neurons. Neuronal development, connectivity and plasticity are dependent on the presence of ER membranes within fine neuronal processes such as spines, axons and dendrites where they are transported along and tethered to microtubules. TAOK2 is critical for dendritic spine development (Ultanir et al., 2014; Yadav et al., 2017), axon elongation and basal dendrite branching (de Anda et al., 2012; Richter et al., 2018). TAOK2 knockout mouse models exhibit cognitive and social-behavioral deficits, and show structural changes in brain size (Richter et al., 2018), mechanisms of which are unknown. Further, de novo mutations in TAOK2 have been found through whole-genome and exome sequencing of patients with autism spectrum disorder (Richter et al., 2018). Our findings provide a hitherto unknown mechanism for kinase mediated control of ER tethering to microtubule cytoskeleton. Elucidating TAOK2 function in specialized cells types such as neurons that respond to physiological stimuli by remodeling ER-microtubules tethering is likely to expand on our current understanding of the importance and dynamics of communication between cellular organelles and the cytoskeleton.

## Acknowledgement

We are grateful for research funding provided by National Institute of Mental Health, R00 MH108648 and R01 MH121674 to SY, the Brain and Behavior Research Foundation’s NARSAD Young Investigator Award (27818) to SY. KN was supported by the NIH pre-doctoral Pharmacological Sciences Training Program T32GM007750. We thank Dan Fong (Nikon) for technical assistance on the Nikon CSU-W1 SoRa superresolution microscope. We are grateful to Brian Beliveau (UW), Cole Trapnell (UW) and Jay Shendure (UW, HHMI) for use of their Nikon CSU-W1 SoRa superresolution microscope funded by HHMI. Thanks to John D. Scott (UW), Ning Zheng (UW, HHMI) and Shao-En Ong (UW) for comments and suggestions on the manuscript.

## Author contribution

All experiments were designed, performed and analyzed by KN and SY, unless otherwise stated. AF generated and characterized the TAOK2 KO lines and cloned the GFP-TAOK2 construct. MB performed molecular biology experiments. Project funding obtained by SY. Manuscript written by KN and SY, and edited by all authors.

## Methods

### Antibodies and Plasmids

Antibodies used in this study are as follows: alpha-Tubulin (Mouse, Sigma, T9026-100UL), GAPDH (Mouse, Invitrogen, MA5-151738), Calreticulin (Mouse, Abcam, ab22683), TAOK2, Rabbit, Sigma, HPA010650), Rtn3a (Rabbit, ProteinTech,12055-2-AP), Acetylated alpha Tubulin (Mouse, Sigma, T6793), GST (Mouse, Invitrogen), GM130 (Mouse, BD Labs, 610822), TAOK2-Cterm (Rabbit), Tom20 (Mouse, Santa Cruz Biotech, sc-17764), Stim1 (Mouse, Santa Cruz Biotech, sc-166840), GFP (Mouse, Roche, 11 814 460 001), Phospho-TAO2 (S181) (Rabbit, R&D Systems, PPS037). Addgene Plasmids used in this study are as follows: mCh-Sec61 beta (#49155), sfGFP-C1 (#54579), ER-mRFP (#62236), EGFP-Sec22b (#101918).

### Molecular Cloning and CRISPR/Cas9 genome editing

Full length human TAOK2 was PCR amplified from pCMV-Sp6-TAOK2 plasmid described previously (Ultanir et al., 2014), and inserted in vector sfGFP-C1 (Addgene #54579) using restriction sites HindIII and MfeI. Domain dissection mutants were subcloned from sfGFP-TAOK2 using restriction enzymes HindIII and MfeI (New England Biolabs). All resultant plasmids were verified by sequencing. GST-TAOK2-(1187-1235) was subcloned from the sfGFP-TAOK2 into the pGEX4T1 vector using sites SalI and NotI. TAOK2 knockout cell line was generated using CRISPR/Cas9 genome editing in HEK293T cells. Four independent guides were designed using Synthego guide design tool (https://www.synthego.com) to target coding exon 2. Two plasmids were made by adding their respective guides into CrisprV2pSpCas9(BB)-2A-Puro (PX459) V2.0 (Addgene Plasmid #62988). Cells were passaged in single cell suspension and plated at 50% confluence. Cultures were then transfected with lipofectamine 2000 reagent (Invitrogen 11668-030) and 2mg of each respective plasmid. Non-Homology End Joining (NHEJ) repair created a deletion around the gRNA cutting site. Cells were selected with Puromycin for 2 days. Genomic DNA was extracted and the region around the cutting site was PCR amplified. Knockout of the gene *TAOK2* was confirmed by Sanger sequencing analysis, and absence of encoded protein was validated using western blot.

### Cell Culture and Maintenance

All experiments were performed in HEK293T cells, which were grown in DMEM media (Thermo Fisher, Gibco) with 10% fetal bovine serum (Axenia BioLogix) and 1% Pen-Strep (Invitrogen). Cells were maintained at 5% CO2 and 37°C and passaged every 3-4 days.

### Immunofluorescence and Western Blotting

Cells were fixed with 4% paraformaldehyde and 4% sucrose for 20 minutes at room temperature, followed by 3 washes with phosphate-buffered saline (PBS). One-hour incubation with blocking buffer (200mM Glycine pH 7.4, 0.25% TritonX-100, 10% Normal Donkey Serum, in PBS) was followed by overnight incubation with primary antibody at a 1:1000 dilution in blocking buffer. After three 5min washes in PBS, cells were incubated with secondary antibody at 1:1000 dilution in blocking buffer for 3hr. Coverslips were washed and then mounted onto slides with FluoromontG. Endogenous TAOK2 staining was performed similarly, except cells were fixed with cold methanol incubated on ice for 20 minutes instead of PFA. Samples for western blot analysis were treated with 4X LDS Sample Buffer (Thermo Fisher) with 125 mM DTT and subsequently heated for 10 minutes at 95C. Samples were electrophoresed on NuPAGE 4-12% Bis-Tris Polyacrylamide gels (Thermo Fisher) with NuPAGE MOPS running buffer (Thermo Fisher). Western blot transfer to ImmobilonP PVDF membrane (Millipore-Sigma) with Transfer Buffer (25mM Tris, 192mM Glycine, 20% (v/v) Methanol, 0.05% SDS) at 100V for 60min. Resultant blot was blocked in 5% milk or BSA blocking buffer, and subjected to primary antibody and HRP conjugated secondary antibody before visualization with Pierce™ ECL Western Blotting Substrate (Thermo Fisher). Western blot images were obtained using the ChemiDoc Imager (BioRad).

### Image Analysis and quantification

MT growth analysis was performed using the ImageJ plugin for manually tracking objects (Manual Tracking). Single frame time lapse image stacks were processed to measure the distance traveled by EB3 comets in each frame. Velocity was calculated by dividing the total distance traveled by the time taken. Curvature was calculated by dividing the minimum linear path (from beginning to end) by the length of the actual distance the EB3 comet traveled. Time paused was calculated by measuring the number of times the EB3 comet did not change coordinates multiplied by time between each frame (2s). Percent time paused is calculated by divided by the total time the comet was tracked.

ER motility was measured from single-z frame image stacks acquired from imaging the ER markers (EGFP-Sec22 or ER-mRFP) using ImageJ. Substacks were created corresponding to frames 6s apart and pixel differences every 6s were calculated using the Stack Difference function to determine a change in fluorescence (ΔF). ΔF was then divided by the mean fluorescence (F) of the earliest time point frame from which it was derived. (i.e., ΔF between frames 3 and 4 would be divided by frame 3) These ΔF/F values were taken for each time point and averaged over 2 minutes to determine ER motility for each cell. To obtain the cumulative pixel difference the substack obtained from using the Stack Difference function was z-projected with the ‘sum slices’ option, and then pseudocolored using the physics LUT. This method to calculate ER motility index is an adaptation from Dong et al. (2018) (Dong et al., 2018) with specified changes.

To assess mitotic defects, four color images were acquired as z-stacks with 0.3micron spacing such that the entire mitotic cell was captured. Mitotic defects were scored manually by visualizing the entire z stack, based on the spindle morphology, ER morphology and chromosomal localization.

### Differential Centrifugation Assay

HEK293T cells were grown to confluence in four 10cm dishes using DMEM media with 10% fetal bovine serum and 1% Pen-Strep. Cells were washed once with Dulbecco’s PBS, collected in ice cold PBS and pelleted by centrifugation at 200g. Pellet was resuspended in 2 mL of homogenization buffer (250mM sucrose, 10mM HEPES, 1mM EDTA, protease inhibitors (Roche), 1mM PMSF, and 1mM DTT) and homogenized with a 25-gauge syringe needle. Homogenate was subsequently spun at 800g to pellet nuclear fraction. Post nuclear supernatant (S1) was diluted in homogenization buffer to split between two 2mL ultracentrifuge tubes. Heavy membrane fraction (P2) was obtained by centrifuging S1 at 27,000g for 30 minutes at 4 °C. Light membrane fraction (P3) was obtained by centrifuging S2 at 100,00g for 30 minutes. P3 was resuspended in either 200uL: homogenization buffer (control), or detergent buffer (1% NP-40, 1% TritonX-100, 0.1% SDS) and incubated on ice for 30 minutes before spinning at 200,000g for 60 minutes at 4 °C. Resultant high speed pellets (P4) were resuspended in 4x sample buffer with 125mM DTT. Resultant supernatants S4 and cytosolic supernatant fraction S3 were precipitated with ice cold 10% trichloroacetic acid by incubating on ice for 15 minutes followed by centrifugation at 21,000g. Precipitates were washed with ice-cold acetone, and pelleted at 21,000g for 5 minutes and resuspended in 4x sample buffer with 125 mM DTT. All samples were run on SDS-PAGE gels and transferred to PVDF membrane for western blot analysis.

### ER Membrane Isolation

HEK293T cells were grown to confluence in four 10cm dishes. Cells were washed once with Dulbecco’s PBS, collected in ice cold PBS and pelleted by centrifugation at 200g. Pellet was resuspended in 1mL of isolation buffer (225mM mannitol, 75mM sucrose, 30mM Tris-HCl pH 7.4, 0.1mM EGTA and protease inhibitors (Roche), and homogenized with a 25G needle at 4°C. The homogenate was subsequently subjected to a series of spins at 4°C, retaining the pellet from each and continuing with supernatant to the next spin. The centrifugation schema was as follows (adapted from Hoyer et al. 2018): 2 spins at 600g for 5 minutes to pellet both cell debris and nuclei (P1 and P2), 3 spins at 7000g for 5 minutes each to pellet mitochondria (P3, P4, and P5), and a 20,000g spin for 20 minutes to pellet the crude ER fraction (P6). The supernatant from the last spin yielded the cytosol, and P6 was washed with isolation buffer devoid of EGTA and subject to a 20,000g spin for 15 minutes at 4°C to re-pellet. P1-6 were resuspended in resuspension buffer (50mM HEPES, 2.5mM MgCl_2_, 200mM KCl, 5% glycerol, 1% TritonX-100). Protein concentrations of resuspended P1-6 and cytosolic fraction were quantified by BCA assay, and subsequently 4X sample buffer with 0.125M DTT was added. This method of crude organellar separation was adapted from Hoyer et al., (2018) (Hoyer et al., 2018). Normalized samples were analyzed by western blot.

### Microscopy

Superresolution imaging was performed using the Nikon-CSU-W1 Spinning Disk equipped with a microlensed SoRa emission disk that achieves Super Resolution by Optical Pixel Reassignment with a xy resolution of 120nm. Images were acquired on an inverted Nikon Eclipse Ti2 microscope (Nikon Instruments) attached to a Yokogawa spinning disk unit (CSU-W1 SoRa, Yokogawa Electric) using a 1.49 100x Apo TIRF oil immersion objective lens. Images were captured by an Andor Sona 4.2B-11 camera using the 2.8x SoRa relay, resulting in an effective pixel size of ∼40 nm. 405, 488, and 561 nm laser lines were used for excitation. All other live and fixed cell imaging was performed on a Nikon Ti2 Eclipse-CSU-X1 confocal spinning disk microscope equipped with four laser lines 405nm, 488nm, 561nm and 670nm and an sCMOS Andor camera for image acquisition. The microscope was caged within the OkoLab environmental control setup enabling temperature and CO_2_ control during live imaging. Imaging was performed using Nikon 1.49 100x Apo 100X or 60X oil objectives. Live imaging for ER motility and EB3 comet velocity was performed on fibronectin coated MatTek dishes (MatTek, P35G-1.5-14-C), and images at a single confocal z frame were captured every 2sec. Fixed cell image acquisition was performed as a z stack of images with z distance of 0.3micron. Viafluor microtubule live imaging dyes (Biotum, #70064, #70063) were used to visualize microtubules during live imaging. Cells were incubated for 30 minutes at 37 degrees C with (1:2000) dye in culture media. Subsequently, dye treated media is replaced with live imaging media with (1:10000) tubulin dye. DNA was stained with NucBlue™ Live ReadyProbes™ Reagent (Hoechst 33342) (Invitrogen), two drops/mL live imaging media incubated for 15 minutes at room temperature before imaging.

### Protein Purification

TAOK2 C-terminal amino acids 1187-1235 were cloned into pGEX4T1 vector and transformed into BL21 *E. coli* to bacterially express the GST-tagged TAOK2 1187-1235. A 25ml starter culture grown from a single colony overnight was used to inoculate 1L culture, which was allowed to grow to an O.D, of 0.6 at 37°C. Protein expression was induced by IPTG at a final concentration of 0.4mM for 18 hours at 18°C. Bacteria were collected by a 15min spin at 5000g, washed with ice cold PBS, and the pellet was resuspended in ice cold lysis buffer (50mM Tris pH 8.0, 5mM EDTA, 150mM NaCl, 20% glycerol, 5mM DTT, protease inhibitors and PMSF). Addition of 4mg lysozyme (Sigma) was followed by 30min incubation with 0.5% TritonX-100 and sonication on ice. The supernatant was collected after a 60min spin at 25,000g, and incubated with prewashed GST beads (Thermo Fisher) for 1 hour. Beads were washed with wash buffer (PBS + 1mM DTT + 0.1% tween 20) followed by wash buffer without detergent. Bound protein was eluted and collected in fractions by glutathione elution buffer at pH8.0 (50mM Tris pH8.0, 250mM KCl, 1mM DTT, 10% glycerol and 30mM glutathione).

### Microtubule co-sedimentation assay

Microtubules were prepared by polymerizing porcine tubulin (Cytoskeleton inc.) in general tubulin buffer (80mM PIPES pH 6.9, 2mM MgCl_2_, 0.4mM EGTA, Roche Protease Inhibitors) in the presence of 1mM GTP for 20 minutes at 35ᵒC and then diluted further. To prevent depolymerization, microtubules were treated with 40 μM Taxol. Microtubules and 5 μg purified protein were incubated at room temperature, and pelleted at 100,000g over a 60% glycerol cushion buffer (80mM PIPES pH 7.0, 1 mM MgCl2, 1 mM EGTA, 60% Glycerol, protease inhibitor). The supernatant (top layer above cushion) and the pellet were removed and treated with 4X Sample Buffer with 250mM DTT and 5% beta-mercaptoethanol. Resultant samples were subject to SDS-PAGE and colloidal Coomassie blue staining (Invitrogen).

### Mitotic cell lysate

HEK293T cells were synchronized by treatment with 1.67μM nocodazole for 12-16 hours. Rounded cells were dislodged by shaking and collected with media. Concurrently, untreated asynchronously growing HEK293T cells were scraped and collected in DPBS. Both tubes of cells were separately pelleted, washed in DPBS, and lysed in HKT buffer (25mM HEPES pH7.2, 150mM KCl, 1% Triton X-100, 1mM DTT, 1 mM EDTA, Protease Inhibitors (Roche, Complete), Halt Phosphatase Inhibitors (Thermo Fisher). Lysate was cleared of cell debris by centrifugation and protein concentrations were determined via BCA assay (Thermo Fisher). Sample Buffer with 125mM DTT was added to equalized amounts of protein and subject to western blot analysis as described above.

### Immunoprecipitation Kinase Assay

HEK293T cells transfected with sfGFP-TAOK2 WT and sfGFP-TAOK2 K57A were lysed with HKT buffer (25mM HEPES pH7.2, 150mM KCl, 1% Triton X-100, 1mM DTT, 1 mM EDTA, Protease Inhibitors (Roche, Complete)). Lysate was precleared with Pierce ProteinG Agarose (Thermo Fisher), and immunoprecipitated with Roche anti-GFP Mouse antibody bound on Pierce protein G agarose. Beads were washed twice with HKT, once for 10 minutes with HKT with 1mM NaCl, and finally washed with HK buffer (25mM HEPES pH 7.2, 150mM KCl, 1mM DTT, 1 mM EDTA, Protease Inhibitors (Roche, Complete EDTA free)). Beads were washed once with the Kinase Buffer (25mM Tris pH 7.5, 10mM MgCl2, 1mM DTT) prior to the in vitro kinase assay. Kinase assay was then performed by incubating with 0.5mM ATP and Halt protease/phosphatase inhibitors (Thermo Fisher) for 45 minutes at 30°C on a shaking heat block. Samples were then subjected to western blot analysis detailed above to detect autophosphorylation of TAOK2 at S181 using the rabbit antibody (R&D Systems, PPS037).

### Statistics

All statistics were performed in GraphPad software Prism9.0. Multiple groups were analyzed using ANOVA, while two group comparisons were made using unpaired t-test unless otherwise stated. Statistically p value less that 0.05 was considered significant. All experiments were done in triplicate, and experimental sample size and p values are indicated with the corresponding figures.

## Supplementary Information

### Supplementary Figure Legend

**Figure S1.**
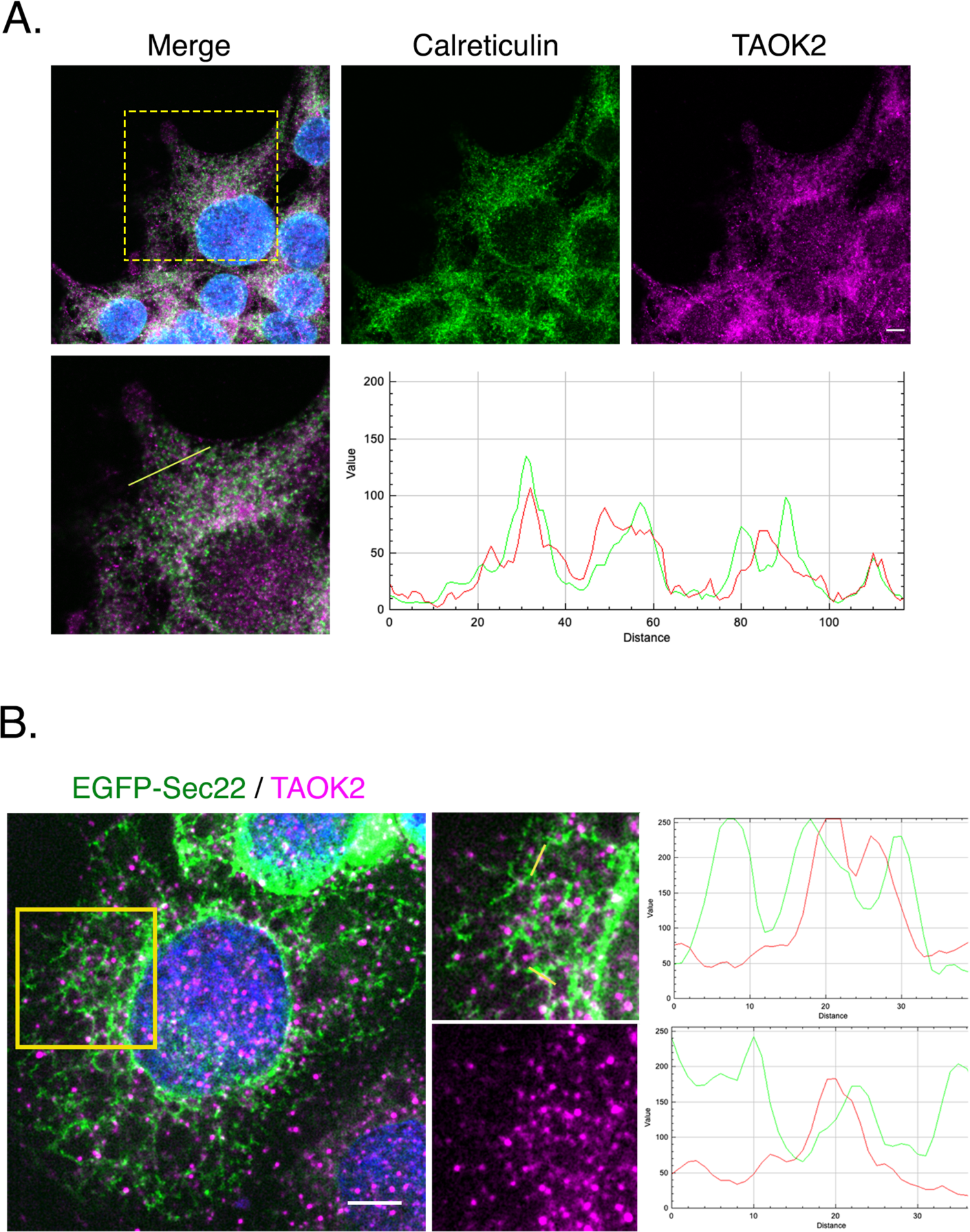
TAOK2 localizes to the ER membrane. (related to main figure 1) (A) Confocal images of HEK293T cells stained with antibodies against calreticulin (green) and TAOK2 (magenta). Higher magnification of peripheral ER (yellow box) is shown in the bottom row. RGB profiler in ImageJ was used to plot the colocalization of calreticulin (green) and TAOK2 (red) fluorescence intensities in the region highlighted by the yellow line. Scale is 5μm. (B) HEK293T cells expressing ER marker EGFP-Sec22b were fixed and stained with antibodies against TAOK2 and DAPI, scale bar is 5μm. Higher magnification of peripheral ER (yellow box) is shown. RGB profiler in ImageJ was used to plot the colocalization of EGFP-Sec22b (green) and TAOK2 (red) fluorescence intensities in distinct subdomains for the region highlighted by the yellow lines. Scale bar is 5μm.

**Figure S2.**
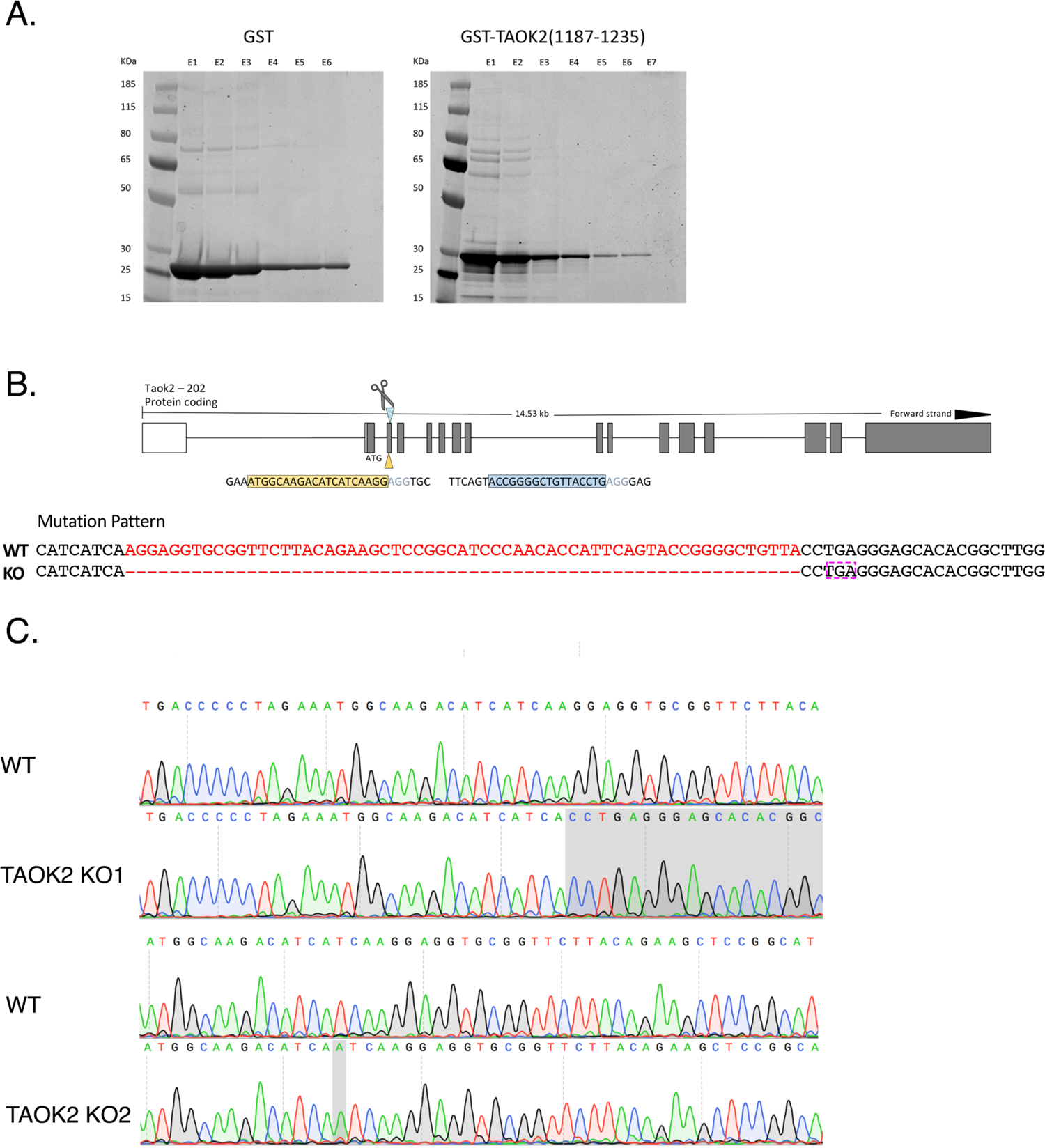
TAOK2 C-terminal tail protein purification and generation of TAOK2 knockout cell line. (related to main figure 2 and 4) (A) Coomassie stained gels showing eluate fractions obtained after affinity purification of GST (left) and GST-TAOK2-C (right) proteins. Fraction E4 was concentrated and used for downstream microtubule binding assays. Molecular weight proteins ladder is shown on the left. (B) Genomic structure of TAOK2 as visualized through the UCSC genomic browser, shows the exons in gray, and region targeted by RNA guides (yellow and blue) is marked by scissors. The resulting deletion caused by genomic editing is shown in red, and premature stop codon is marked by the magenta box. (C) Sequence peaks show the result of DNA sequencing in wildtype and two separate KO cell lines performed after PCR of the surrounding genomic region. The change in sequence is shown in gray.

**Figure S3.**
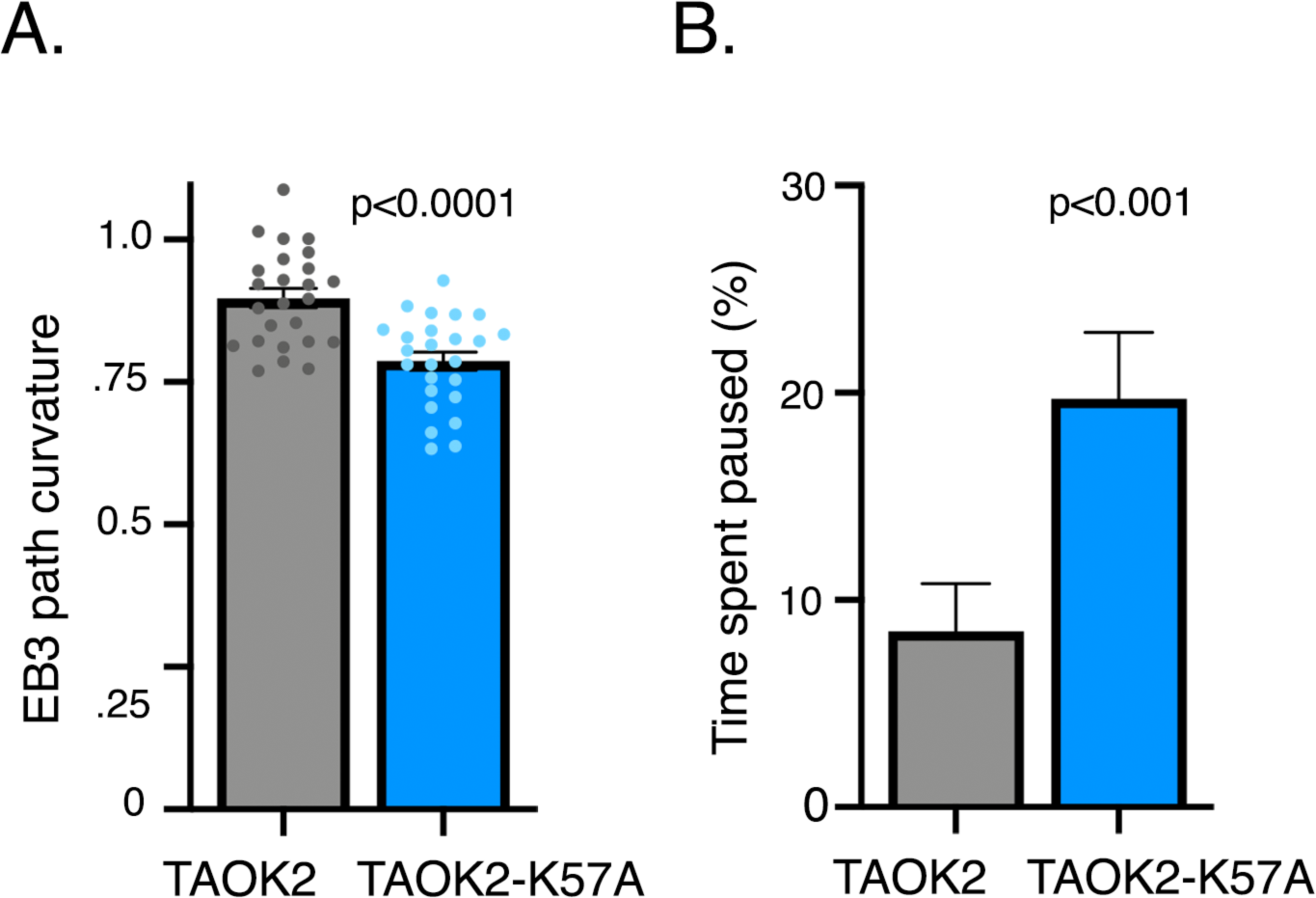
Effect of kinase dead TAOK2 K57A on microtubule dynamics. (related to main figure 6) (A) Microtubule directional persistence calculated as a fraction of perpendicular distance between start/end points and the length of the actual path taken is plotted for cells expressing TAOK2-WT and TAOK2-K57A. Values indicate mean ± S.E.M., n=10 cells with at least 5 comet paths measured per cell, two tailed t-test. (B) Percent total time spent by EB3 comet pausing (no growth) is plotted for TAOK2-WT and TAOK2-K57A expressing cells. Values indicate mean ± S.E.M., n=10 cells with at least 5 comet paths measured per cell, two tailed t-test.

### Movie Legends

**Supplementary Movie 1. (related to Figure 1)** TAOK2 is an ER protein. Combined time-lapse image stacks obtained on CSU-W1 SoRa superresolution confocal microscope show representative HEK293T cell transfected with GFP-TAOK2 (yellow) and ER-mRFP (magenta) to visualize the dynamics of TAOK2 on ER membranes. Image stacks were acquired every 2 sec. Movie compiled at 8 frames per sec (fps). Scale bar = 1μm.

**Supplementary Movie 2. (related to Figure 3)** TAOK2 is an ER-MT tether. Combined time-lapse image stacks show representative HEK293T cell transfected with GFP-TAOK2 and ER-mRFP. Cells were incubated with a MT binding dye (405nm excitation) for 30min prior to acquiring images on a confocal microscope. Triple channel images were acquired at a single focal plane every 2s to simultaneously visualize TAOK2 (yellow), ER (magenta), and microtubules (cyan). Note localization of TAOK2 punctae at points of contact between ER and microtubules. Movie compiled at 8fps. Scale bar = 2μm.

**Supplementary Movie 3. (related to Figure 3)** Aberrant unregulated TAOK2 tethering disrupts ER-MT dynamics. Combined time-lapse image stacks show representative HEK293T cell transfected with GFP-TAOK2(1146-1235), ER-mRFP (magenta) and stained with microtubule dye (cyan) to visualize the dynamics of ER membranes and MT growth. Complete absence of ER motility in presence of TAOK2 tether can be compared to normal ER membrane dynamics in an adjacent cell not transfected with GFP-TAOK2 (1145-1235) construct. Image stacks were acquired every 2 sec. Movie compiled at 8 frames per sec (fps).

**Supplementary Movie 4. (related to Figure 4)** Combined time lapse-image stacks show representative wildtype and TAOK2 knockout cell transfected with EGFP-Sec22b (cyan) and mCherry-EB3 (red) to visualize the dynamics of ER membranes and MT growth. Image stacks were acquired every 2 sec. Movie compiled at 8 frames per sec (fps).

**Supplementary Movie 5. (related to Figure 6)** Combined time-lapse image stacks show representative HEK293T cell transfected with TAOK2-WT, EGFP-Sec22b (magenta) and mCherry-EB3 (blue-green) to visualize the dynamics of ER membranes and MT growth. Image stacks were acquired every 2 sec. Movie compiled at 8 frames per sec (fps).

**Supplementary Movie 6. (related to Figure 6)** Combined time-lapse image stacks show representative HEK293T cell transfected with TAOK2-K57A, EGFP-Sec22b (magenta) and mCherry-EB3 (blue-green) to visualize the dynamics of ER membranes and MT growth. Image stacks were acquired every 2 sec. Movie compiled at 8 frames per sec (fps).

## Notes

### Competing Interest Statement

The authors have declared no competing interest.

